# MET Receptor Tyrosine Kinase Regulates Vagal Laryngeal Motor Neuron Development and Lifespan Ultrasonic Vocal Communication

**DOI:** 10.1101/2020.05.13.093617

**Authors:** Anna K. Kamitakahara, Ramin Ali Marandi Ghoddousi, Alexandra L. Lanjewar, Valerie M. Magalong, Hsiao-Huei Wu, Pat Levitt

## Abstract

The vagal motor nucleus ambiguus (nAmb) innervates the intrinsic muscles of the larynx, providing direct motor control over vocal production in humans and rodents. Here, we demonstrate that early developmental signaling through the MET receptor tyrosine kinase (MET) is required for proper formation of the nAmb. Embryonic deletion of *Met* in the developing brainstem resulted in a loss of one-third of motor neurons in the nAmb. While the remaining neurons were able to establish connections with target muscles in the larynx, advanced signal processing analyses revealed severe deficits in ultrasonic vocalization in early postnatal life. Abnormal vocalization patterns persisted into adulthood in the majority of mice tested. Interestingly, 28% of adult mice recovered the ability to vocalize demonstrating heterogeneity in circuit restitution. Together, the data establish MET as a factor necessary for development of a specific subset of neurons in the nAmb required for normal ultrasonic vocalization.

## Introduction

The generation of functionally distinct motor circuits is dependent on developmental exposure to a defined set of signaling cues. The precise identity of these instructive signals and their roles in distinguishing each of the discrete motor neuron pools is central to our understanding of neurodevelopmental processes, disorders, and neural control of all of the muscle groups in the body.

The research focus historically has been on spinal motor neuron development (Jessell, 2000; Ladle et al., 2007; Purves, 1988). Yet, among the most functionally diverse populations of motor neurons are those of the medullary vagal motor nuclei, which consist of neurons located in the nucleus ambiguus (nAmb) and dorsal motor nucleus of the vagus (DMV). These nuclei control a vast number of functionally distinct muscular targets, including those important for swallowing, vocalization, heart rate, and gastrointestinal motility. The process of delineating these neurons developmentally begins following early brainstem patterning events in the neural plate and tube, coordinated by retinoic acid exposure, and Hox gene induction (Gavalas et al., 1998; Linville et al., 2004; Philippidou and Dasen, 2013). Between embryonic day (E)9.5 and E10.5 in mice, vagal sensory, motor, and other central and peripheral autonomic neurons are then derived from Phox2b expressing precursor cells (Dauger et al., 2003; Pattyn et al., 1999; Pierce, 1973). The fate of vagal motor neurons is established subsequently through Islet-1 expression (Dubreuil et al., 2000; Ericson et al., 1992), followed by Hox5 expression and late retinoic acid exposure driving rostral-caudal patterning of vagal motor neurons (Barsh et al., 2017; Gavalas et al., 1998; Isabella et al., 2020; Linville et al., 2004). What remains unknown are the mechanisms through which functional diversity of the vagal motor neurons is attained, and the factors that guide the development of individual motor neuron subclasses.

Recent work from our laboratory and others have demonstrated that prenatally, subsets of developing vagal motor neurons express the Met receptor tyrosine kinase (MET) (Isabella et al., 2020; Kamitakahara et al., 2017; Wu and Levitt, 2013), positioning it as a candidate for distinguishing motor neuron types. MET is a pleiotropic receptor, known to be important in synapse maturation and critical period development in the cerebral cortex (Chen et al., 2020; Ma and Qiu, 2019; Xie et al., 2016) and for the differentiation of several different motor neuron pools in the brainstem and spinal cord (Caton et al., 2000; Ebens et al., 1996; Tallafuss and Eisen, 2008; Wong et al., 1997). For example, upon binding its only known ligand, hepatocyte growth factor (HGF), MET confers neuronal survival in developing spinal motor neurons innervating the pectoralis minor muscle (Lamballe et al., 2011). By contrast, in adjacent limb-innervating motor neurons, MET is dispensable for neuronal survival but instead stimulates axonal elaboration (Lamballe et al., 2011). The distinct effects of MET signaling in these different aspects of neurodevelopment (i.e. neuronal survival vs. axonal growth) may be the result of the combination of other trophic factors that are co-expressed with MET in discrete motor neuron pools (Isabella et al., 2020; Schaller et al., 2017; Wong et al., 1997). Additionally, coordination of the timing between MET expression in motor neurons and HGF expression in target innervation sites further ensures proper innervation patterns (Isabella et al., 2020; Lamballe et al., 2011). Similar to spinal motor neurons, a recent study in zebrafish suggests that MET signaling is involved in axon growth and guidance of vagal motor neurons innervating the pharyngeal arches (Isabella et al., 2020). Mammalian species exhibit considerable differences from the fishes in vagal anatomy and pharyngeal arch-derived structures, as mammals form a pharynx and larynx instead of gill arches. However, analogous to zebrafish, HGF is expressed embryonically in the developing airways and esophagus in mice (Kamitakahara et al., 2017). This suggests that in mammals, MET may serve as a signal instructing the development of esophageal and laryngeal motor neurons, which are located in the nAmb.

Here, we used a conditional knockout approach to delete *Met* from vagal motor neurons expressing Cre recombinase under the control of the Phox2b promoter (*Phox2b*^*cre*^*;Met*^*fx/fx*^ mice) to determine the functional role of MET signaling during development, and the potential enduring impact of developmental disruptions. Using this transgenic mouse model, we examined the development of the nAmb and the structural abnormalities that arise from the loss of MET. Furthermore, we analyzed the effects of the loss of MET signaling on neuromuscular junction formation at innervation sites in the larynx and upper esophagus. Lastly, we used advanced signal processing methods to determine the physiological effects of conditional deletion of *Met* on organ function, focusing on the larynx, which is responsible for production of ultrasonic communication. Heterogeneous, but enduring vocalization disturbances in the conditional knockout reveals a novel role for MET signaling in vagal motor neuron specialization and function.

## Results

### Transgenic model for conditional loss of MET in vagal motor neurons

To examine the function of the MET receptor tyrosine kinase (MET) in motor neurons in the Nucleus Ambiguus (nAmb), we employed a Cre-Lox strategy to conditionally delete MET from developing vagal neurons. *Phox2b*^*cre*^ mice were crossed to the *Met*^*fx/fx*^ transgenic mouse line. In this model, Cre-mediated recombination facilitates excision of exon 16 of the *Met* allele (the functional ATP binding site), such that only truncated, non-functional protein is transcribed (Huh et al., 2004). The Cre-dependent reporter tdTomato (tdTom) was further used to identify all vagal populations following functional deletion of MET, providing a method to identify and count nAmb motor neurons in the developing and adult medulla. As early as E9.5, tdTom+ neurons were present in rhombomeres 7/8, the region from which vagal motor neurons originate (Lumsden and Keynes, 1989), indicating that recombination and deletion of MET occurs early, just following cell birth (Supplemental Figure 1).

To confirm loss of MET from vagal motor neurons, immunohistochemistry was used to examine protein expression in vagal axonal projections. While no antibodies are available to exclusively detect the portion of the receptor transcribed by exon 16, work from our lab and others have demonstrated the signaling incompetence and subsequent degradation of MET protein following Cre-mediated recombination in *Met*^*fx/fx*^ mice (Huh et al., 2004; Judson et al., 2010; Peng et al., 2016). In agreement with these results, in control *Phox2b*^*cre*^*;Met*^+*/*+^ mice, abundant MET immunolabeling could be visualized in both the nAmb and axonal projections of developing vagal motor neurons on E14.5 (Figure 1A, B, E, and E’). By contrast, in conditional knockout *Phox2b*^*cre*^*;Met*^*fx/fx*^ mice, MET immunolabeling could only be detected in the nAmb, but not in axonal projections (Figure 1C, D, F, and F’) suggesting that truncated MET protein is degraded in the cell soma and not transported to innervation targets. Using these transgenic mouse lines, we were able to test functional changes in vagal motor development resulting from the loss of MET.

**Figure 1.**
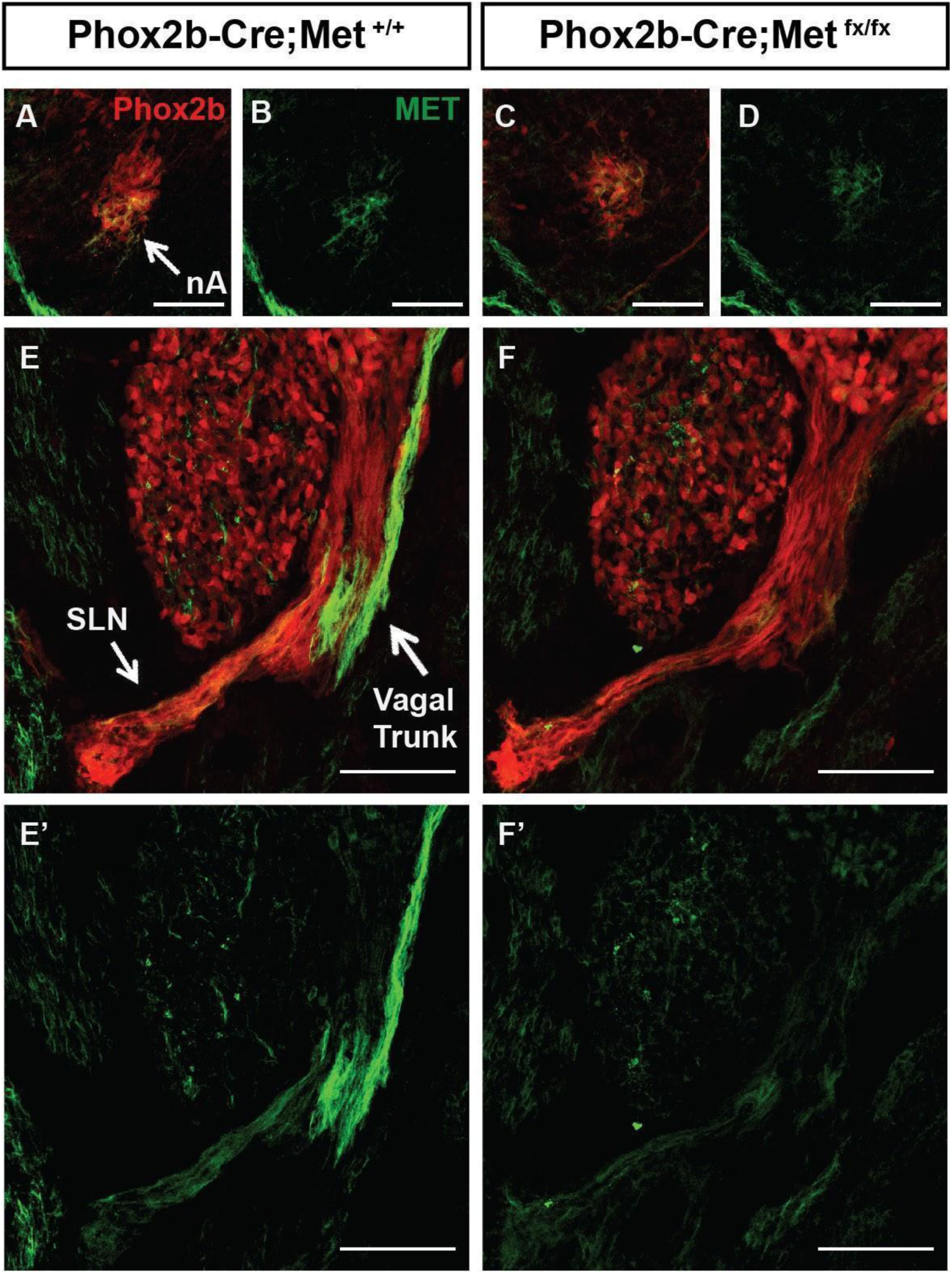
Conditional deletion of *Met* results in early embryonic loss of MET protein in vagal motor neuron axons. (A, B) Colocalization of MET protein (green) with Phox2b (tdTomato, red) in the nAmb of the *Phox2b*^*cre*^*;Met*^+*/*+^ mouse on E14.5. (C, D) Colocalization of MET protein lacking the functional ATP activation site (green) with Phox2b (tdTomato, red) in the nAmb of *Phox2b*^*cre*^*;Met*^*fx/fx*^ mouse on E14.5. (E,E’) Expression of MET protein (green) in the tdTom+ (red) neuronal fibers of the vagal trunk and the superior laryngeal nerve (SLN) in a *Phox2b*^*cre*^*;Met*^+*/*+^ mouse. (F,F’) Minimal expression of MET protein in the neuronal fibers of the vagal trunk and the superior laryngeal nerve (SLN) in a *Phox2b*^*cre*^*;Met*^*fx/fx*^ mouse. All scalebars = 100μm. n = 4 mice per group. The brightness and contrast of each channel was adjusted separately for visualization purposes.

### MET is required for formation of the nAmb

To examine a potential role for MET in the development of the nAmb, we first performed analysis of brainstem sections collected from *Phox2b*^*cre*^*;Met*^*fx/fx*^*;tdTom* mice at several time points, beginning in embryonic life through adulthood. On E14.5, a decrease in the size of the developing nAmb in *Phox2b*^*cre*^*;Met*^*fx/fx*^*;tdTom* samples was evident (Figure 2A). While the size of the nucleus was reduced, there were no apparent, ectopically located tdTom neurons in the medulla. Quantification of the number of neurons in the nAmb at this time point revealed a significant 23.3% decrease in the number of tdTom+ neurons in *Phox2b*^*cre*^*;Met*^*fx/fx*^*;tdTom* mice compared to *Phox2b*^*cre*^*;Met*^+*/*+^*;tdTom* controls (Figure 2B). To determine whether this loss in neuron number is maintained later in life, tdTom+ cell counts were quantified in early postnatal and adult brainstem samples. The early decrease in nAmb cell number observed prenatally grew to 36.4% and 37.8% decreases in tdTom+ neurons on P7 and P60, respectively (Figure 2C-F), demonstrating that MET is required during embryonic and postnatal development for sustaining a normal number of nAmb neurons.

**Figure 2.**
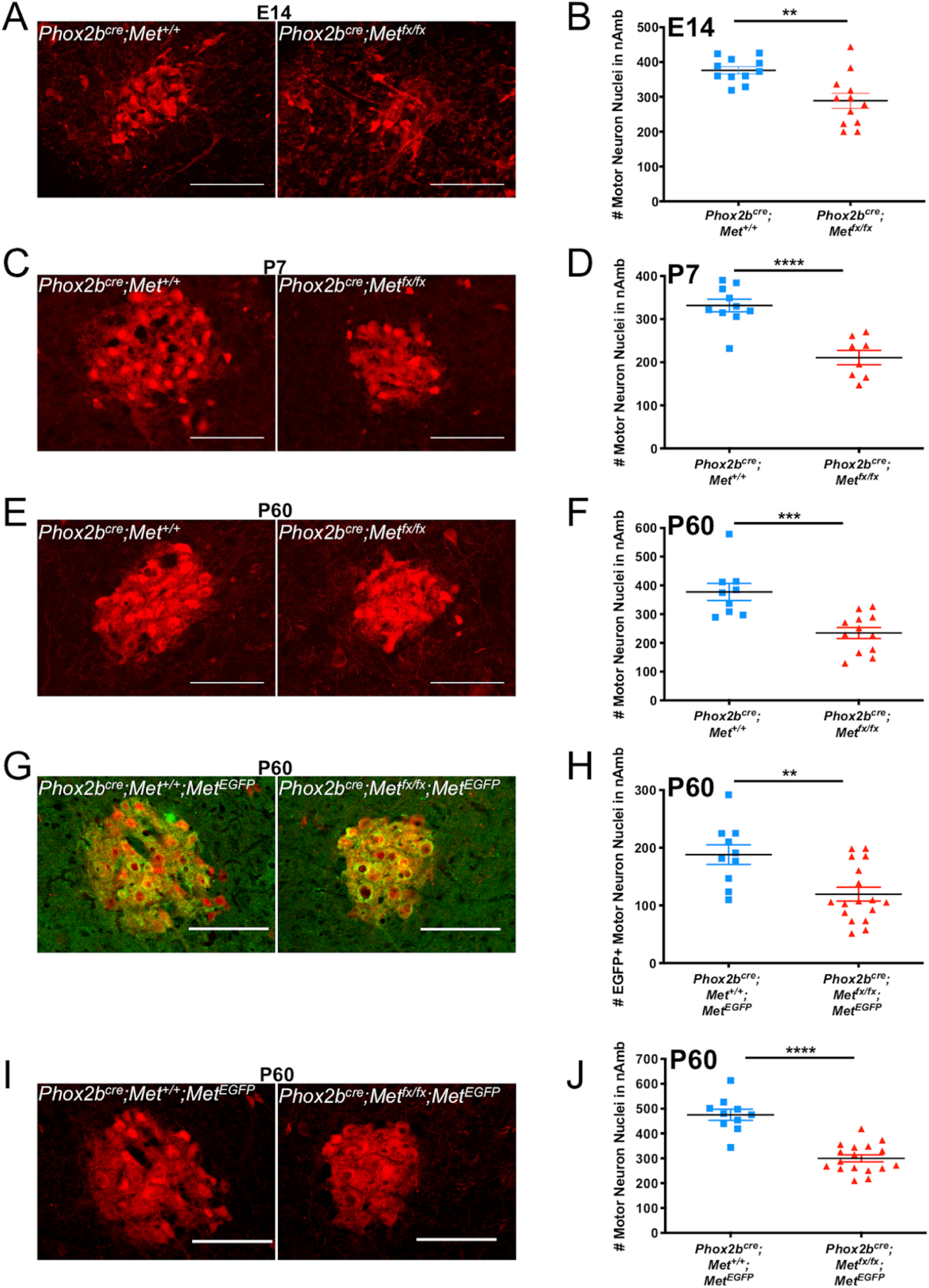
Loss of vagal motor neurons in the nAmb following conditional deletion of MET. (A) Representative images of the nAmb from control and *Phox2b*^*cre*^*;Met*^*fx/fx*^*;tdTom* mice on E14.5. (B) tdTom+ cell counts from control and *Phox2b*^*cre*^*;Met*^*fx/fx*^*;tdTom* mice on E14.5. n = 11 *Phox2b*^*cre*^*;Met*^+*/*+^*;tdTom*, 12 *Phox2b*^*cre*^*;Met*^*fx/fx*^*;tdTom*. (C) Representative images of the nAmb of control and *Phox2b*^*cre*^*;Met*^*fx/fx*^*;tdTom* mice on P7. (D) tdTom+ cell counts from control and *Phox2b*^*cre*^*;Met*^*fx/fx*^*;tdTom* mice on P7. n = 10 *Phox2b*^*cre*^*;Met*^+*/*+^*;tdTom*, 8 *Phox2b*^*cre*^*;Met*^*fx/fx*^*;tdTom*. (E) Representative images of the nAmb of control and *Phox2b*^*cre*^*;Met*^*fx/fx*^*;tdTom* mice on P60. (F) tdTom+ cell counts from control and *Phox2b*^*cre*^*;Met*^*fx/fx*^*;tdTom* mice on P60. n = 11 *Phox2b*^*cre*^*;Met*^+*/*+^*;tdTom*, 12 *Phox2b*^*cre*^*;Met*^*fx/fx*^*;tdTom*. (G) Representative images of the GFP+ (green) motor neurons (red) in nAmb of control and *Phox2b*^*cre*^*;Met*^*fx/fx*^*;tdTom;Met*^*EGFP*^ mice on P60. (H) EGFP+ cell counts from control and *Phox2b*^*cre*^*;Met*^*fx/fx*^*;tdTom;Met*^*EGFP*^ mice on P60. n = 10 *Phox2b*^*cre*^*;Met*^+*/*+^*;tdTom;Met*^*EGFP*^, 17 *Phox2b*^*cre*^*;Met*^*fx/fx*^*;tdTom:Met*^*EGFP*^. (I) Representative images of the motor neurons (red) in nAmb of control and *Phox2b*^*cre*^*;Met*^*fx/fx*^*;tdTom;Met*^*EGFP*^ mice on P60. (J) tdTom+ cell counts from control and *Phox2b*^*cre*^*;Met*^*fx/fx*^*;tdTom;Met*^*EGFP*^ mice on P60. n = 10 *Phox2b*^*cre*^*;Met*^+*/*+^*;tdTom;Met*^*EGFP*^, 17 *Phox2b*^*cre*^*;Met*^*fx/fx*^*;tdTom:Met*^*EGFP*^. All scalebars = 100μm. The brightness and contrast of each channel was adjusted separately for visualization purposes. Some images were cropped to center the nAmb in the image plane.

Previously published work from our laboratory suggests that only a subset of the neurons in the nAmb express MET. These neurons are primarily located in the compact and loose formation of the nAmb during development (Kamitakahara et al., 2017). To determine whether the loss of neurons observed in the nAmb corresponds to the subpopulation of the nucleus expressing MET, *Phox2b*^*cre*^*;Met*^*fx/fx*^*;tdTom* mice were further crossed with mice expressing enhanced green fluorescent protein (EGFP) under the control of the MET promoter (*Met*^*EGFP*^). In control *Phox2b*^*cre*^*;Met*^+*/*+^*;tdTom;Met*^*EGFP*^ mice, this permitted visualization of neurons expressing MET, and facilitated comparison in conditionally deleted *Phox2b*^*cre*^*;Met*^*fx/fx*^*;tdTom;Met*^*EGFP*^ mice, in which the *Met*^*EGFP*^ reporter labeled neurons producing non-functional MET protein (Figure 2G). Cell counts in the nAmb on P60 revealed a 36.4% decrease in the number of EGFP+ neurons (Figure 2H), consistent with the absence of only a subpopulation of *Met*^*EGFP*^ expressing neurons in the nAmb following functional loss of MET. Likewise, no apparent loss in EGFP+ neurons was observed in the DMV of a small subset of samples analyzed (data not shown), providing further evidence of a selective impact on a subset of vagal motor neurons located in nAmb.

To validate that the same loss of nAmb motor neurons occurs in the *Phox2b*^*cre*^*;Met*^*fx/fx*^*;tdTom; Met*^*EGFP*^ mouse line as seen in the *Phox2b*^*cre*^*;Met*^*fx/fx*^*;tdTom* mouse line, total tdTom+ neurons in the nAmb were quantified at P60 and reveal a 36.9% decrease in the tdTom+ neurons, as expected (Figure 2I-J). Furthermore, within each genotype, no sex differences were observed in the number of EGFP+ or tdTom+ neurons at any of the postnatal ages tested (Supplemental Figure 2).

### Laryngeal and esophageal muscles are innervated following nAmb cell loss

To determine whether the loss of a subset of nAmb neurons results in the loss of innervation to its peripheral targets in the esophagus and larynx, these tissues containing tdTom+ axons were stained with fluorescently tagged alpha-bungarotoxin (*α*BT) to label acetylcholine receptor (AChR) clusters at the neuromuscular junction. The percentage of *α*BT closely apposed to tdTom+ axons was then quantified in control and *Phox2b*^*cre*^*;Met*^*fx/fx*^*;tdTom* mice on P7. In the esophagus, nearly all of the *α*BT labeled AChR clusters were apposed by tdTomato+ axons in both genotypes (Figure 3A-C), suggesting that esophageal neuromuscular junctions form without MET. Similarly, in all of the intrinsic muscles in the larynx responsible for vocal production that were assayed (thyroarytenoid, TA; lateral cricoarytenoid, LCA; cricothyroid, CT; posterior cricoarytenoid, PCA; and alar portion of the thyroarytenoid, AlarTA), nearly all of the *α*BT labelled AChR clusters were contacted by tdTomato+ axons in both genotypes, suggesting that laryngeal neuromuscular junctions form without MET (Figure 3D-L).

**Figure 3.**
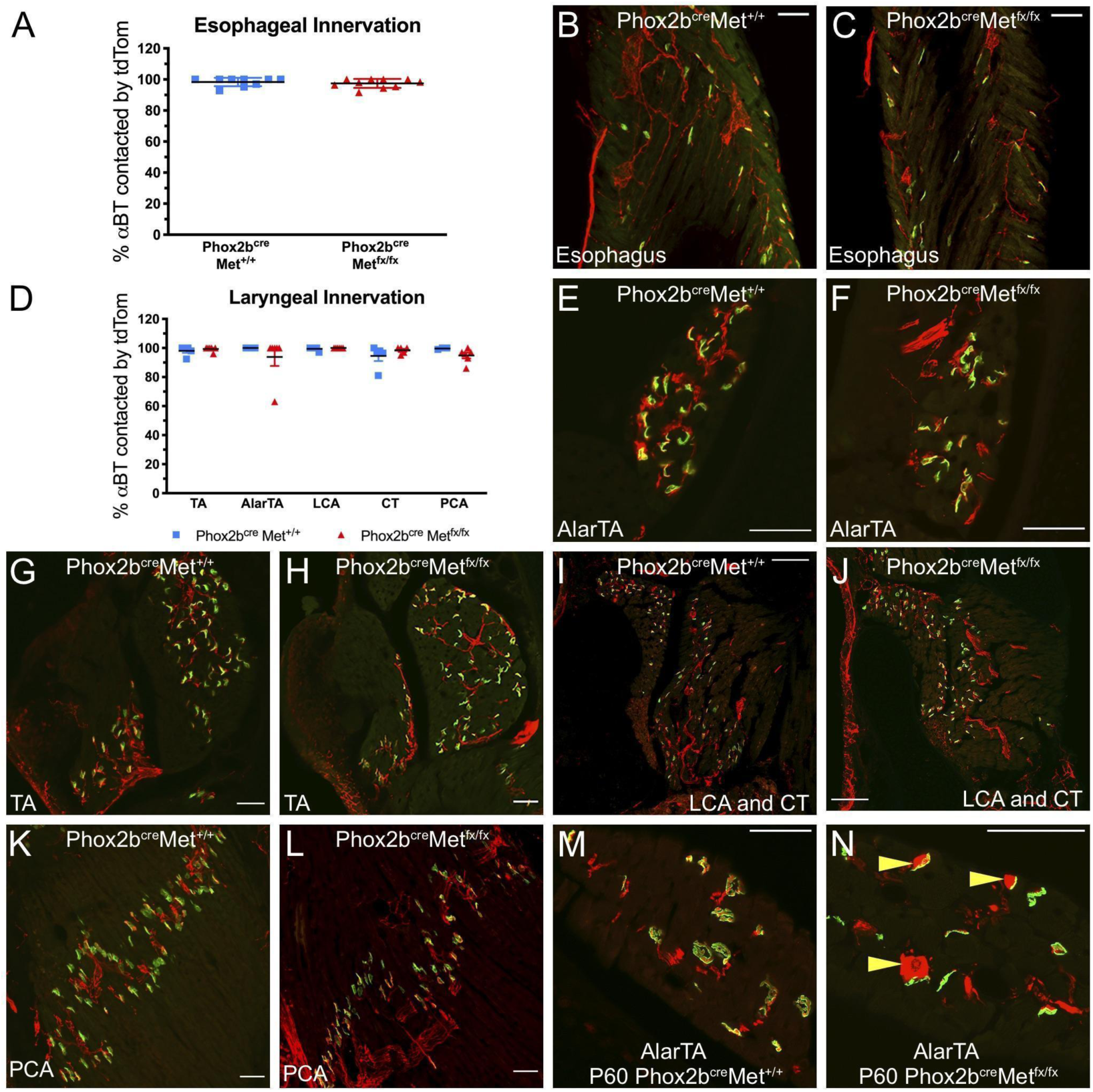
*α*BT labelled AChR clusters are innervated in esophageal and laryngeal muscles is at P7 following MET deletion. (A) Quantification of esophageal innervation in control and *Phox2b*^*cre*^*;Met*^*fx/fx*^*;tdTom* mice on P7. n = 9 *Phox2b*^*cre*^*;Met*^+*/*+^*;tdTom*, 10 *Phox2b*^*cre*^*;Met*^*fx/fx*^*;tdTom*. (B,C) Representative images of tdTom+ axons (red) innervating *α*BT labelled AChR clusters (green) in the esophagus of control and *Phox2b*^*cre*^*;Met*^*fx/fx*^*;tdTom* mice on P7. (D) Quantification of laryngeal innervation in control and *Phox2b*^*cre*^*;Met*^*fx/fx*^*;tdTom* mice on P7. n = 3-5 *Phox2b*^*cre*^*;Met*^+*/*+^*;tdTom*, 5-6 *Phox2b*^*cre*^*;Met*^*fx/fx*^*;tdTom*. (E,F) Representative images of tdTom+ axons (red) innervating *α*BT labelled AChR clusters (green) in the alar thyroarytenoid (AlarTA) muscle of control and *Phox2b*^*cre*^*;Met*^*fx/fx*^*;tdTom* mice on P7. (G,H) Representative images of tdTom+ axons (red) innervating *α*BT labelled AChR clusters (green) in the thyroarytenoid muscle (TA) of control and *Phox2b*^*cre*^*;Met*^*fx/fx*^*;tdTom* mice on P7. (I,J) Representative images of tdTom+ axons (red) innervating *α*BT labelled AChR clusters (green) in the lateral cricoarytenoid (LCA) and cricothyroid (CT) muscles of control and *Phox2b*^*cre*^*;Met*^*fx/fx*^*;tdTom* mice on P7. (K,L) Representative images of tdTom+ axons (red) innervating *α*BT labelled AChR clusters (green) in the posterior cricoarytenoid (PCA) muscle of control and *Phox2b*^*cre*^*;Met*^*fx/fx*^*;tdTom* mice on P7. (M) Typical neuromuscular junction morphology observed in the AlarTA in control mice on P60. (N) Swollen presynaptic tdTom+ axons (red) innervating *α*BT labelled AChR clusters (green) in the AlarTA of *Phox2b*^*cre*^*;Met*^*fx/fx*^*;tdTom* mouse on P60. All scalebars = 50μm The brightness and contrast of each channel was adjusted separately for visualization purposes.

Neuromuscular junction maintenance was also assessed in esophageal and laryngeal samples from *Phox2b*^*cre*^*;Met*^*fx/fx*^*;tdTom* mice on P60. At this time point, almost all *α*BT labelled AChR clusters were contacted by tdTom+ axons in the esophagus and intrinsic laryngeal muscles of both genotypes (data not shown). However, within the AlarTA, an unusual, swollen presynaptic profile was observed in 57% (4 out of 7) of the *Phox2b*^*cre*^*;Met*^*fx/fx*^*;tdTom* mice, and in only 20% (1 out of 5) of control littermates (Figure 3N). While the exact mechanisms by which USVs are produced remain a contested area of research (Mahrt et al., 2016; Riede et al., 2017; Roberts, 1975), one hypothesis suggests that movement of the alar cartilage may be critical for rodent vocalization through an edge-tone whistle mechanism (Riede et al., 2017). The AlarTA is the only muscle connected to the alar cartilage. Thus, the morphological phenotype observed in the AlarTA provided initial evidence that despite the formation of neuromuscular junctions by the remaining nAmb neurons, neuronal function might otherwise be altered.

### Ultrasonic vocalization is impaired in early postnatal life following loss of MET

The activity of vagal laryngeal motor neurons in the nAmb is required for rodent ultrasonic vocalization (USV) (Nunez et al., 1985; Wetzel et al., 1980; Yajima et al., 1982). Furthermore, despite the finding that *α*BT labeled AChR clusters receive innervation in *Phox2b*^*cre*^*;Met*^*fx/fx*^ pups, it is unknown whether those connections are functional, or require MET for coordinated activity. To test whether vagal motor output is functionally altered following loss of MET, USVs evoked by isolation of mouse pups from the nest were recorded on P7. Following removal from the nest, control *Phox2b*^*cre*^*;Met*^+*/*+^ pups evoked robust USV calls over the course of a 5-minute recording period. Both the number and duration of calls were similar between Cre negative and *Phox2b*^*cre*^*;Met*^+*/*+^ pups, demonstrating that Cre expression alone in this transgenic model has no effect on vocalization (Figure 4A and B). By contrast, *Phox2b*^*cre*^*;Met*^*fx/fx*^ pups exhibited severely reduced numbers of calls and decreased call duration (Figure 4A and B), indicating that the loss of MET in vagal motor neurons impairs USV production. Remarkably, at this age, there was nearly uniform penetrance of the USV phenotype. To test whether the loss of isolation-evoked USVs is due to a motor deficit to call, or perhaps a reduced motivation to call, short 15-second recordings were made following a brief tail pinch stimulus to directly elicit vocalization. While both Cre negative and *Phox2b*^*cre*^*;Met*^+*/*+^ pups evoked many complex USVs following the tail pinch, *Phox2b*^*cre*^*;Met*^*fx/fx*^ pups made few calls, if any, and those emitted were very short in duration (data not shown). These results suggest that pups lacking MET are unable to make complex calls, and that the neurons expressing MET in the nAmb are critical for laryngeal motor neuron function.

**Figure 4.**
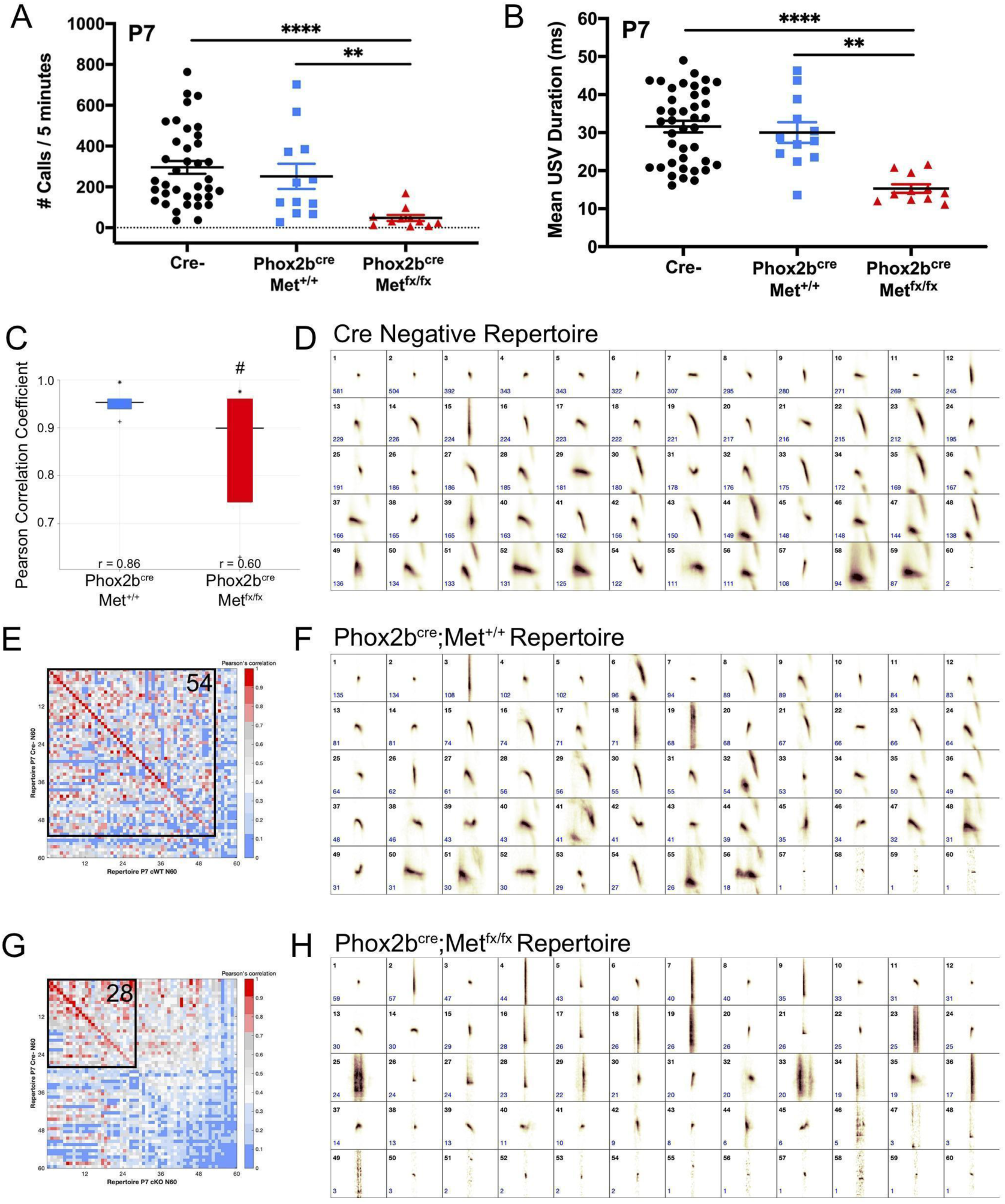
Deletion of MET results in severely impaired ultrasonic vocalization evident in early postnatal life. (A) Quantification of the number of isolation-evoked USVs made over the 5-minute recording period on P7. n = 37 Cre Negative, n = 12 *Phox2b*^*cre*^*;Met*^+*/*+^*;tdTom*, n = 11 *Phox2b*^*cre*^*;Met*^*fx/fx*^*;tdTom*. (B) Quantification of the duration of isolation-evoked USVs made over the 5-minute recording period on P7. n = 37 Cre Negative, n = 12 *Phox2b*^*cre*^*;Met*^+*/*+^*;tdTom*, n = 11 *Phox2b*^*cre*^*;Met*^*fx/fx*^*;tdTom*. (C) MUPET boxplot comparing the Cre negative repertoire to the *Phox2b*^*cre*^*;Met*^+*/*+^ repertoire (blue) or *Phox2b*^*cre*^*;Met*^*fx/fx*^ repertoire (red). Similarity of the top 5% most frequently used RUs in the Cre negative repertoire indicated by *. Similarity of the top 25% most frequently used RUs in the Cre negative repertoire indicated by top of the box. Similarity of the top 50% most frequently used RUs in the Cre negative repertoire indicated by the horizontal line. Similarity of the top 75% most frequently used RUs in the Cre negative repertoire indicated by the bottom of the box. Similarity of the top 95% most frequently used RUs in the Cre negative repertoire indicated by +. r values below boxes indicate the overall Pearson correlation coefficient for the entire repertoire. # indicates overall Pearson’s r values that are significantly different from the Cre negative repertoire. (D) Each repertoire unit (RU) in the Cre negative repertoire, displayed in the order of frequency of use. (E,G) Pearson correlation matrices comparing each of the Cre negative RUs (y-axis) to RUs in the *Phox2b*^*cre*^*;Met*^+*/*+^ repertoire (x-axis, E) or *Phox2b*^*cre*^*;Met*^*fx/fx*^ repertoire (x-axis, G), ordered from most to least similar in shape. Warmer colors indicate higher Pearson correlation, cooler colors indicate lower Pearson correlation. Boxed area shows the number of RUs with Pearson correlations above 0.7, with the corresponding number of RUs indicated in the upper right corner. (E) Each repertoire unit (RU) in the *Phox2b*^*cre*^*;Met*^+*/*+^ repertoire, displayed in the order of frequency of use. (G) Each repertoire unit (RU) in the *Phox2b*^*cre*^*;Met*^*fx/fx*^ repertoire, displayed in the order of frequency of use.

Rodent USV repertoires are composed of a diverse compliment of syllable shapes used to generate strain, sex, and context specific vocal communication across development and in adulthood (Arriaga et al., 2012; Grimsley et al., 2011; Holy and Guo, 2005; Segbroeck et al., 2017). Given that loss of MET results in a major reduction in the number and duration of calls made by *Phox2b*^*cre*^*;Met*^*fx/fx*^ mice postnatally, we hypothesized that the shape of syllables emitted might also be altered, reflecting impairments in the ability to vocalize. To analyze differences in USV repertoires between genotypes, Mouse Ultrasonic Profile ExTraction (MUPET) software was used to perform unsupervised signal processing of isolation-evoked USVs recorded on P7 (Segbroeck et al., 2017). Within each genotype, all recorded syllables were clustered into repertoires composed of 60 repertoire units (RUs). Each RU represents the syllable centroid, or average of syllable shapes in that RU cluster. The Cre negative repertoire was composed of a number of distinct RUs (short simple, frequency jumps, complex harmonics, etc.) (Figure 4D). The shape of each of these RUs in the Cre negative repertoire was then statistically compared to RUs clustered in *Phox2b*^*cre*^*;Met*^+*/*+^ and *Phox2b*^*cre*^*;Met*^*fx/fx*^ repertoires (Figure 5F and H). For this analysis, MUPET generates a matrix of individual Pearson correlations between pairwise comparisons of RUs in each repertoire. The matrix is color coded such that warmer colors correspond to more highly similar RUs. Fifty-four out of 60 RUs had Pearson correlations greater than 0.7 in the *Phox2b*^*cre*^*;Met*^+*/*+^ repertoire (colored red to pink across the matrix diagonal in Figure 4E), demonstrating that the shape of the syllables in these repertoires are highly similar to one another. In contrast, the *Phox2b*^*cre*^*;Met*^*fx/fx*^ repertoire was primarily composed of very short syllables, and a straight vertical shape often associated with either clicking sounds or noise artifacts. Only 28 out of 60 RUs in the *Phox2b*^*cre*^*;Met*^*fx/fx*^ repertoire had Pearson correlations greater than 0.7 when compared to the Cre negative repertoire (Figure 4G), and were primarily made up of very short simple shapes, suggesting that the subset of nAmb neurons expressing MET is critical for generating the diversity and complexity of syllables in the vocal repertoire.

**Figure 5.**
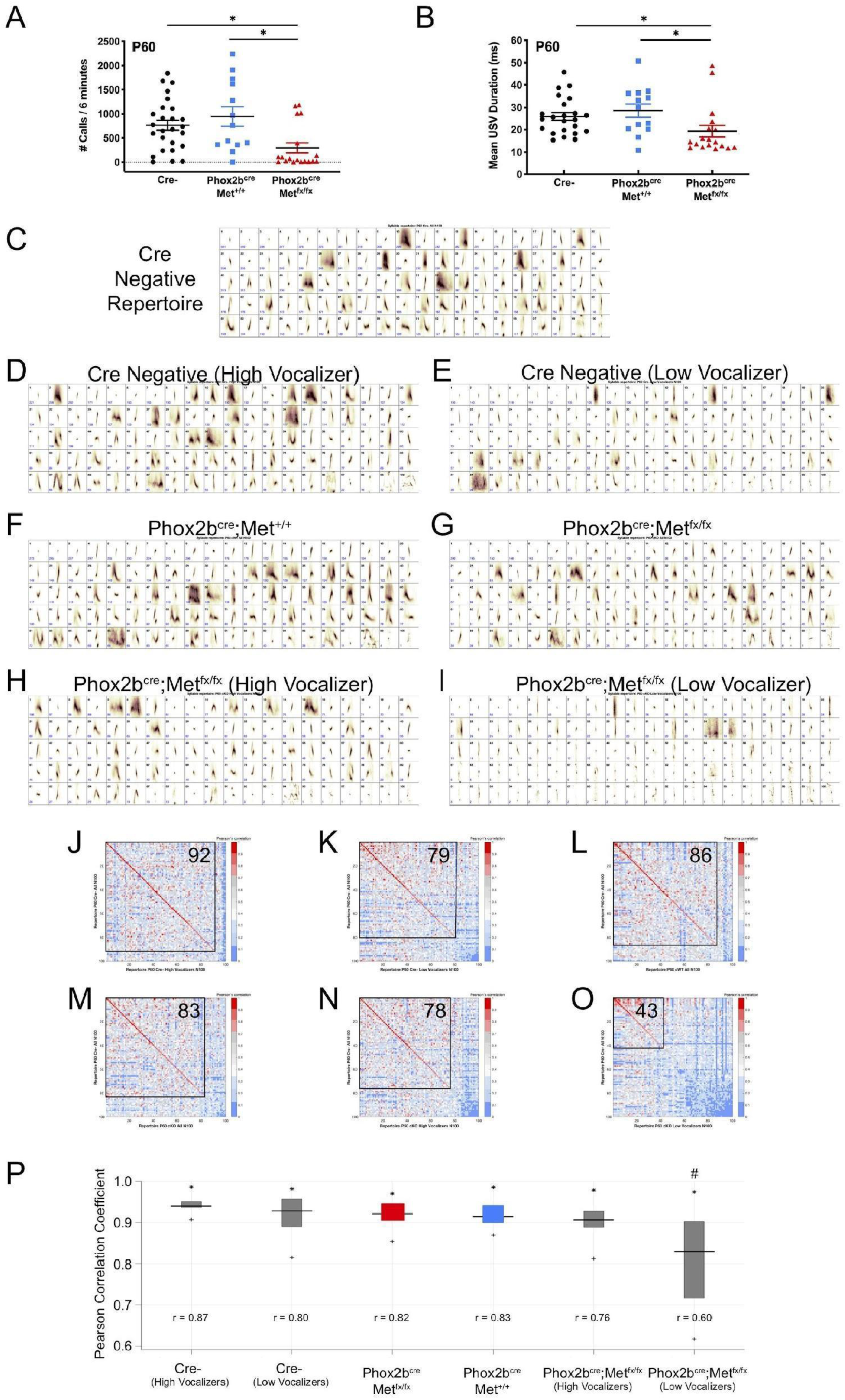
Recovery of vocalization deficits observed in a subset of mice lacking MET in vagal motor neurons. (A) Quantification of the number of USVs made by males paired with females over the 6-minute recording period on P60 during a direct social interaction task. n = 26 Cre Negative, 13 *Phox2b*^*cre*^*;Met*^+*/*+^*;tdTom*, 18 *Phox2b*^*cre*^*;Met*^*fx/fx*^*;tdTom* (B) Quantification of the duration of USVs made by males over the 6-minute recording period on P60. n = 26 Cre Negative, 13 *Phox2b*^*cre*^*;Met*^+*/*+^*;tdTom*, 18 *Phox2b*^*cre*^*;Met*^*fx/fx*^*;tdTom* (C) Each repertoire unit (RU) in the Cre-repertoire, displayed in the order of frequency of use. (D) Each repertoire unit (RU) in the Cre-high vocalizer (HV) repertoire, displayed in the order of frequency of use. Repertoire generated from 13 Cre Negative recordings with the highest number of calls. (E) Each repertoire unit (RU) in the Cre-low vocalizer (LV) repertoire, displayed in the order of frequency of use. Repertoire generated from 13 Cre Negative recordings with the lowest number of calls. (F) Each repertoire unit (RU) in the *Phox2b*^*cre*^*;Met*^+*/*+^ repertoire, displayed in the order of frequency of use. (G) Each repertoire unit (RU) in the *Phox2b*^*cre*^*;Met*^*fx/fx*^ repertoire, displayed in the order of frequency of use. (H) Each repertoire unit (RU) in the *Phox2b*^*cre*^*;Met*^*fx/fx*^ high vocalizer (HV) repertoire, displayed in the order of frequency of use. Repertoire generated from 5 *Phox2b*^*cre*^*;Met*^*fx/fx*^*;tdTom* recordings with the highest number of calls. (I) Each repertoire unit (RU) in the *Phox2b*^*cre*^*;Met*^*fx/fx*^ low vocalizer (LV) repertoire, displayed in the order of frequency of use. Repertoire generated from 13 *Phox2b*^*cre*^*;Met*^*fx/fx*^*;tdTom* recordings with the lowest number of calls. (J, K, L, M, N, O) Pearson correlation matrices comparing each of the Cre-RUs (y-axis) to RUs in the Cre-high vocalizer (HV) repertoire (x-axis, J), Cre-low vocalizer (LV) repertoire (x-axis, K), *Phox2b*^*cre*^*;Met*^+*/*+^ repertoire (x-axis, L), *Phox2b*^*cre*^*;Met*^*fx/fx*^ repertoire (x-axis, M), *Phox2b*^*cre*^*;Met*^*fx/fx*^ high vocalizer (HV) repertoire (x-axis, N), or *Phox2b*^*cre*^*;Met*^*fx/fx*^ low vocalizer repertoire (x-axis, O), ordered from most to least similar in shape. Warmer colors indicate higher Pearson correlation, cooler colors indicate lower Pearson correlation. Boxed area shows the number of RUs with Pearson correlations above 0.7, with the corresponding number of RUs indicated in the upper right corner. (P) MUPET boxplot comparing the Cre negative repertoire to each group. Similarity of the top 5% most frequently used RUs in the Cre negative repertoire indicated by *. Similarity of the top 25% most frequently used RUs in the Cre negative repertoire indicated by top of the box. Similarity of the top 50% most frequently used RUs in the Cre negative repertoire indicated by the horizontal line. Similarity of the top 75% most frequently used RUs in the Cre negative repertoire indicated by the bottom of the box. Similarity of the top 95% most frequently used RUs in the Cre negative repertoire indicated by +. r values below boxes indicate the overall Pearson correlation coefficient for the entire repertoire. # indicates overall Pearson’s r values that are significantly different from the Cre negative repertoire.

In addition to comparing individual RUs, groups of the most frequently used RUs in the Cre negative repertoire were compared to RUs in the *Phox2b*^*cre*^*;Met*^+*/*+^ and *Phox2b*^*cre*^*;Met*^*fx/fx*^ repertoires. As the top 5% most frequently used RUs across all genotypes tended to be short simple syllables, Pearson correlation coefficients comparing both the *Phox2b*^*cre*^*;Met*^+*/*+^ and *Phox2b*^*cre*^*;Met*^*fx/fx*^ were high: 1.00 and 0.98, respectively (Figure 4C). To compare the entire Cre negative repertoire to the entire *Phox2b*^*cre*^*;Met*^+*/*+^ and *Phox2b*^*cre*^*;Met*^*fx/fx*^ repertoires, Pearson correlation coefficients were calculated for 100% of used syllables. While the *Phox2b*^*cre*^*;Met*^+*/*+^ repertoire was highly similar to the Cre negative repertoire, with an overall Pearson’s r of 0.86, the *Phox2b*^*cre*^*;Met*^*fx/fx*^ repertoire was significantly different from the Cre negative repertoire, with an overall Pearson’s r of 0.60 (Figure 4C). This difference in overall repertoire similarity suggests that MET expression by the nAmb is required for the generation of the full vocal repertoire produced by P7 mouse pups.

### USVs are impaired in the majority of male Phox2b^cre^;Met^fx/fx^ mice in adulthood

To determine whether the inability to emit USVs is sustained into adulthood, separate cohorts of Cre negative, *Phox2b*^*cre*^*;Met*^+*/*+^ and *Phox2b*^*cre*^*;Met*^*fx/fx*^ male mice were examined at P60 using a direct social interaction task with a female C57BL/6J conspecific. Only males were analyzed in this paradigm, as females did not produce USVs in sufficient numbers at this age to distinguish from baseline (data not shown). Both Cre negative and *Phox2b*^*cre*^*;Met*^+*/*+^ males averaged several hundred calls over the course of the 6-minute recording period (Figure 5A). By contrast, *Phox2b*^*cre*^*;Met*^*fx/fx*^ male mice produced significantly fewer USVs over that same time period. Furthermore, average USV duration was significantly reduced in mutant mice (Figure 5A and B). Intriguingly, the responses of *Phox2b*^*cre*^*;Met*^*fx/fx*^ male mice appeared to follow two distinct patterns: a majority of mice tested that vocalized very little, using only short simple syllables (low vocalizers), and a smaller group of mice that vocalized fairly robustly, able to use more complex syllables (high vocalizers). This indicates that while the majority of adult *Phox2b*^*cre*^*;Met*^*fx/fx*^ mice exhibited a very limited ability to vocalize, a small subset (5/18; 27.8%) appeared to regain laryngeal function sometime after P7.

To compare USV repertoire deficits observed in *Phox2b*^*cre*^*;Met*^*fx/fx*^ mice, and to determine whether impairments observed in early postnatal life are sustained into adulthood, MUPET was used to examine syllable repertoires in recordings from P60 males paired with females in the direct social interaction task. Within each genotype, all recorded syllables were clustered into 100 RUs, based on modeling parameters. The Cre negative repertoire was composed of a diverse array of simple and complex syllable shapes (Figure 5C). Similarly, based on Pearson correlations above 0.7, both the *Phox2b*^*cre*^*;Met*^+*/*+^ and *Phox2b*^*cre*^*;Met*^*fx/fx*^ repertoire were composed of syllables with 86/100 and 83/100 similarity to the Cre negative repertoire, respectively (Figure 5F, G, L, M). Given observations that the USV recordings from *Phox2b*^*cre*^*;Met*^*fx/fx*^ mice at P60 followed one of two patterns (low vocalizers or high vocalizers), repertoires of these sub-sampled groups were compared. USV recordings from *Phox2b*^*cre*^*;Met*^*fx/fx*^ mice were grouped into either low (n=13) or high vocalizing (n=5) groups split by the mean number of USVs in that *genotype*. USV recordings from Cre negative mice were also grouped into either low or high vocalizing groups split by the mean number of USVs in that genotype. When each of these subgroups was compared to the Cre negative repertoire, it was found that both Cre negative high and low vocalizers had high Pearson correlations for syllable shape when compared to the entire Cre negative group (Figure 5D, E, J, K). This shows that even in control mice that vocalized less frequently, the syllables produced are very similar to those of mice that vocalized more frequently. Similarly, 78/100 syllables from the five *Phox2b*^*cre*^*;Met*^*fx/fx*^ high vocalizers exhibited high Pearson correlations for syllable shape (Figure 5H and N), indicating that both the frequency and quality of vocalizations in this small group of mice were normal, even in the absence of *Met*. In contrast, for the *Phox2b*^*cre*^*;Met*^*fx/fx*^ low vocalizer group, only 43/100 syllables had Pearson correlations above 0.7 (Figure 5I and O), revealing that not only were vocalizations less frequent, but in addition, the repertoires of these mice also were distinct from the Cre negative controls. This is consistent with the apparently “normal” repertoire of the undivided *Phox2b*^*cre*^*;Met*^*fx/fx*^ group being driven by the high vocalizers. Similar to recordings at P7, the *Phox2b*^*cre*^*;Met*^*fx/fx*^ low vocalizer recordings were composed primarily of very short syllables and a straight vertical shape that is often associated with either a clicking sound or noise artifact.

Groups of the most frequently used RUs in the Cre negative repertoire also were compared to RUs in the other genotypes. Similar to observations at P7, the top 5% most frequently used RUs across all genotypes at P60 were short simple syllables, and thus were highly similar between all groups (Figure 5P). To compare the entire Cre negative repertoire to entire repertoires from other genotypes, Pearson correlation coefficients were calculated for 100% of used syllables. Both Cre negative high and low vocalizers had high Pearson correlations for syllable shape when compared to the entire Cre-group, 0.87 and 0.80, respectively (Figure 5P). Similarly, *Phox2b*^*cre*^*;Met*^+*/*+^, *Phox2b*^*cre*^*;Met*^*fx/fx*^, and *Phox2b*^*cre*^*;Met*^*fx/fx*^ high vocalizers had high overall Pearson correlations coefficients when the entire repertoire was compared (Figure 5P). However, *Phox2b*^*cre*^*;Met*^*fx/fx*^ low vocalizers had a significantly lower Pearson’s r value (Figure 5P). This comprehensive repertoire analysis further suggests that the seemingly “normal” repertoire of the undivided *Phox2b*^*cre*^*;Met*^*fx/fx*^ group is likely driven by the few high vocalizers. Together, the data reveal that in the majority of mice tested, deficits in ultrasonic vocal communication are sustained into adulthood.

## Discussion

The development of functional motor circuits is dependent upon the timely exposure of neurons (and their responsiveness) to specific morphogens, motogens, trophic factors, and axon guidance cues. Differing combinations of each provide a means for subtype-specific developmental maturation of motor neurons that innervate distinct muscle groups. In the current study, we identify MET as a critical neurodevelopmental signal that is necessary for establishing normal anatomy and function of vagal motor circuits. Specifically, the present data demonstrate that the loss of MET signaling results in a significant decrease in the number of vagal efferent motor neurons in the nAmb and major deficits in the ability to communicate using ultrasonic vocalizations. The quantitative morphological studies also reveal that like spinal motor neurons, MET+ nAmb motor neurons exhibit differences in receptor dependency during development, indicating a conserved ontogenetic mechanism across distinct efferent motor neuron types. In addition, differential MET-dependence subdivides nAmb motor neurons even further. The functional analyses highlight an unexpected capacity for the remaining nAmb motor neurons to target and form neuromuscular junctions with all of the intrinsic laryngeal muscles in the absence of MET. Despite this maintained structural connection, the profound vocalization abnormalities observed suggest that expression of MET by nAmb motor neurons is necessary for USV production.

### Developmental loss of a subpopulation of nAmb neurons

The MET receptor mediates a pleiotropic neurodevelopmental signal with well-established roles in neuronal proliferation and survival, chemoattraction, and dendritic elaboration and plasticity (Avetisyan et al., 2015; Caton et al., 2000; Ebens et al., 1996; Gutierrez et al., 2004; Isabella et al., 2020; Judson et al., 2010; Lamballe et al., 2011; Lim and Walikonis, 2008; Maina et al., 1998; Nakamura et al., 1989; Peng et al., 2016). Consistent with functioning developmentally, conditional deletion of *Met* in vagal motor neurons resulted in a statistically significant reduction in the number of neurons in the nAmb, detected just a few days after neurogenesis begins embryonically. Examination of the medulla postnatally did not reveal an obvious accumulation of ectopically located tdTom-labeled neurons, suggesting that the reduction of neurons in nAmb was not due to major defects in cell migration. Though we cannot completely exclude that some cells in the ventricular zone express MET, cell labeling is most evident in the mantle zone of the medulla (Figure S1). This is consistent with our studies of the developing telencephalon (Judson et al., 2010) and those in the spinal cord that found the majority of MET expression to be localized to non-proliferating cells (Ebens et al., 1996). We suggest that the prenatal phenotype is consistent with required MET signaling for survival of a subset of nAmb motor neurons.

Virtually all neuronal populations are produced in excess during development, and then undergo a period of naturally occurring cell death (NOCD), eliminating up to half of these neurons between E11.5 and P1 (Hamburger and Levi-Montalcini, 1949; Hollyday and Hamburger, 1976; Hollyday et al., 1977; Oppenheim, 1991; Purves, 1988; White et al., 1998). The classical model of NOCD posits that competition for a limited supply of trophic factors produced by the target muscle serves as one of the central mechanisms through which apoptosis is regulated. This effect is well demonstrated in developing chick spinal motor neurons, where removal of a limb bud in early life, which thereby removes muscle-derived trophic support, results in survival of only a small fraction of lumbar motor neurons on the side of the amputation (Hamburger and Levi-Montalcini, 1949). Conversely, transplantation of a supernumerary limb results in survival of additional motor neurons on the same side as the transplant, due to the increased trophic signal provided by the extra limb (Hollyday and Hamburger, 1976; Hollyday et al., 1977). Further work has identified various trophic factors, including HGF, which protect specific neuronal subpopulations during NOCD. For example, there is a dose-dependent survival of lumbar motor neurons with increasing concentrations of supplemented HGF or chick muscle-derived extract (Ebens et al., 1996; Novak et al., 2000). While HGF supports the survival of lumbar neurons, it has little effect on brachial or thoracic motor neuron survival (Novak et al., 2000), demonstrating the specificity of trophic factor matching. Previous studies from our laboratory demonstrate expression of HGF at nAmb target innervation sites in the muscles of the esophagus and larynx at E13.5 and E15.5 (Kamitakahara et al., 2017), positioning it as a likely candidate mediating survival in this subpopulation of nAmb neurons.

Caspase-dependent cleavage of the five C-terminal amino acids from the MET receptor is thought to mechanistically drive apoptosis in the absence of receptor activation (Duplaquet et al., 2020; Foveau et al., 2007; Ma et al., 2014). It is important to note that in the mouse model used in the current study, the floxed portion of the receptor is well outside this p40MET C-terminal site (Huh et al., 2004), and does not itself experimentally induce an apoptotic response. This is confirmed in other independently published studies (Avetisyan et al., 2015; Judson et al., 2010; Lamballe et al., 2011), as well as in the current study, where conditional deletion of MET in vagal motor neurons results only in a partial loss of EGFP+ neurons in the nAmb of *Met*^*EGFP*^ mice, and no loss is observed in other regions in the dorsal motor nucleus of the vagus.

While loss of MET signaling results in a reduction of neurons in the nAmb, we demonstrate that a majority subset of EGFP+ neurons in *Met*^*EGFP*^ mice are resilient to the deletion of *Met*, surviving even with the loss of MET-HGF signaling. This is consistent with the partial loss of subsets of neuronal populations observed in other developmental trophic factor interactions. For example, GDNF-deficient mice exhibit a loss of approximately one quarter of spinal lumbar neurons despite expression of the GDNF receptors, Ret and Gfra1, in all spinal motor neurons (Moore et al., 1996; Oppenheim et al., 2000). Interestingly, the surviving spinal lumbar neurons also expressed, and even upregulated expression of the receptor Gfra2, suggesting their survival could be supported by other GDNF family members, such as neurturin (Oppenheim et al., 2000). Similarly, in addition to expression of MET, the nAmb has been reported to express receptors for a number of other trophic factors, including p75NTR, TrkA (Gibbs and Pfaff, 1994; Koh et al., 1989), TrkB (Liu and Wong-Riley, 2013), TrkC (Helke et al., 1998), leukemia inhibitory factor receptor (Li et al., 1995), ciliary neurotrophic factor receptor alpha (MacLennan et al., 1996), gp130 (Nakashima et al., 1999), and Gfra1 (Mikaels et al., 2000). Here it is possible that the remaining *Met*^*EGFP*^ neurons that do not express MET are able to maintain their survival through expression or upregulation of receptors for other trophic factors.

### Maintenance of muscle connectivity

Despite the loss of more than a third of the neurons in the nAmb of *Phox2b*^*cre*^*;Met*^*fx/fx*^ mice, innervation at esophageal and laryngeal neuromuscular junctions appeared largely normal, though we cannot eliminate the possibility of subtle changes in the number of motor endplates. Moreover, based on the functional analyses of vocalizations during development and in the adult, a subset of these connections may not be able to mediate normal functions, and/or segregate appropriately into motor pools. Thus, aberrant motor neuron targeting or non-functional sprouting could underlie what appears to be relatively normal structural innervation in the laryngeal muscle groups.

Each of the muscles of the larynx control different properties of vocal production. The thyroarytenoid muscles contract to adduct and shorten the vocal folds, while the cricothyroid muscles tense the vocal folds (Ludlow, 2005; Riede, 2013). The posterior cricoarytenoids abduct the vocal folds, moving them away from the midline (as when breathing), while the lateral cricoarytenoids adduct the vocal folds, bringing them together (as when vocalizing) (Ludlow, 2005). One possibility, therefore, is that disorganized innervation or innervation of muscles with opposing actions by a common neuron results in vocal deficits similar to those observed in *Phox2b*^*cre*^*;Met*^*fx/fx*^ mice.

There are several relevant examples in humans that reflect inappropriate laryngeal innervation that can be heterogeneous. A neurological condition, synkinesis, involves contraction of one muscle occurring in concert with additional muscle groups that typically do not contract together. For example, during thyroid surgery, some patients experience complications causing recurrent laryngeal nerve injury. Subsequent nonselective reinnervation of laryngeal muscle groups can lead to synkinesis (Crumley, 1979, 2000; Flint et al., 1991). Several studies have examined neurotrophin therapy as a potential method of encouraging growth and appropriate reinnervation following recurrent laryngeal nerve injury. Expression of NGF, BDNF, GDNF, and Netrin-1 in the posterior cricoarytenoid have been shown to initially increase, and later decline following recurrent laryngeal nerve injury modeled in rats (Araki et al., 2006; Hernandez-Morato et al., 2014; Hernandez-Morato et al., 2016; WANG et al., 2016). In other preclinical models, NT-3 showed promising results for increasing reinnervation of the posterior cricoarytenoid muscle (Kingham et al., 2007).

Synkinesis also can arise developmentally as demonstrated in mouse models of strabismus where chemokine receptor deletion has been shown to result in improper ocular muscle innervation by trigeminal motor neurons (Whitman et al., 2018). To our knowledge, no studies have specifically demonstrated laryngeal synkinesis of a developmental origin. However, HGF exhibits survival, growth promoting, and chemoattractive properties on the neuronal projections innervating the developing brachial arches (Caton et al., 2000; Ebens et al., 1996; Isabella et al., 2020). Therefore, the loss of MET-expressing laryngeal motor neurons in the nAmb during development could allow for non-selective reinnervation by other motor neuron types in *Phox2b*^*cre*^*;Met*^*fx/fx*^ mice and account for the observed vocal impairments. In addition, girls with Rett Syndrome (RTT) may exhibit disrupted vagal motor circuit function, reflected in atypical feeding, swallowing, and vocalization observed in some infants who are diagnosed later with RTT (Einspieler and Marschik, 2019). This is of interest here, because *MET* transcription is attenuated by mutations in *MECP2*, and *MET* expression is nearly undetectable in postmortem brain samples of girls diagnosed with Rett Syndrome compared to matched controls (Plummer et al., 2013).

### Deficits in ultrasonic vocalization in early development and adulthood

The most dramatic phenotype observed in *Phox2b*^*cre*^*;Met*^*fx/fx*^ mice is the severe disruption of ultrasonic vocal communication. This is most pronounced developmentally when compared to control pups, *Phox2b*^*cre*^*;Met*^*fx/fx*^ pups made few, if any, isolation-evoked USVs during the 5-minute recording period. Furthermore, the application of advanced MUPET syllable repertoire analysis demonstrated that *Phox2b*^*cre*^*;Met*^*fx/fx*^ pups only produce short, simple calls that are likely easier to produce than the more complex long duration calls that are made by the *Phox2b*^*cre*^*;Met*^+*/*+^ pups. Experiments in which the recurrent laryngeal nerves are unilaterally or bilaterally transected demonstrate the importance of this connectivity for vocal fold movement and ultrasonic vocalization (Calub et al., 2018; Nunez et al., 1985; Pomerantz et al., 1983; Roberts, 1975; Wetzel et al., 1980). Despite strong evidence that recurrent laryngeal projections from the nAmb are responsible for USV production in rodents, the exact acoustic mechanisms through which USVs are produced remain an active area of investigation (Mahrt et al., 2016; Riede et al., 2017; Roberts, 1975). The current study narrows the mechanism of action, as the vagal motor circuits responsible for USVs require MET signaling during development for their survival or proper function.

Similar to studies done in rodents, vocal impairments or dysphonia can occur in humans that experience complications during thyroid surgery that result in recurrent laryngeal nerve paralysis (Hartl et al., 2005). In experimental models of recurrent laryngeal nerve injury, synkinesis is observed in 66% to 88% of cases, while in the remainder of cases, reinnervation appears to be normally reinstated (Flint et al., 1991). This reflects a considerable amount of heterogeneity in injury and reinnervation models. Using our rodent model of developmental deletion of MET, we also observe variability in responses, with approximately 28% of *Phox2b*^*cre*^*;Met*^*fx/fx*^ mice able to vocalize normally in adulthood. This is consistent with some level of capacity for plasticity to recover from an initial loss of function developmentally. Further studies, focusing on the minority of mice that exhibit normal adult vocalization, may yield important information for developing new strategies for promoting functional plasticity in individuals with developmentally or surgically induced deficits in laryngeal function and vocal ability.

## Acknowledgements

The research was supported by the Simms/Mann Chair in Developmental Neurogenetics, the WM Keck Chair in Neurogenetics (P.L.), the The Saban Research Institute Program in Developmental Neuroscience and Neurogenetics, The Saban Research Institute’s Research Career Development Fellowship (A.K.K.), and the USC Neuroscience Predoctoral Training Grant T32GM113859 (ALL).

## Author Contributions

Conceptualization, A.K.K., R.A.M.G., H.H.W., and P.L.; Methodology, A.K.K.; Formal Analysis, A.K.K., R.A.M.G, A.L.L., and V.M.M.; Investigation, A.K.K., R.A.M.G, A.L.L., and V.M.M.; Writing – Original Draft, A.K.K.; Writing – Review and Editing, A.K.K., R.A.M.G, A.L.L., V.M.M., H.H.W., and P.L.; Visualization, A.K.K., R.A.M.G, A.L.L., and V.M.M.; Supervision, A.K.K. and P.L.; Project Administration, A.K.K. and P.L.; Funding Acquisition, A.K.K., P.L., and A.L.L.

## STAR*Methods

### Experimental Model

#### Animals

Animal care and experimental procedures were performed in accordance with the Institutional Animal Care and Use Committee of The Saban Research Institute, Children’s Hospital Los Angeles. Mice were housed in the vivarium on a 13:11 hour light:dark cycle (lights on at 06:00 hours, lights off at 19:00 hours) at 22°C with ad libitum access to a standard chow diet (PicoLab Rodent Diet 20, #5053, St. Louis, MO).

The *Phox2b*^*cre*^ and cre-dependent TdTomato reporter lines (*TdTomato*) were obtained from The Jackson Laboratory (B6(Cg)-Tg(Phox2b-cre)3Jke/J, stock 016223, RRID:IMSR_JAX:016223 and B6.Cg-Gt(ROSA)26Sortm14(CAG-tdTomato)Hze/J, stock 007914; Ai14, RRID:IMSR_JAX:007914, respectively). The *Met*^*fx*^ mouse line, in which exon 16 of the *Met* allele is flanked by loxP sites, was shared by the laboratory of Dr. Snorri S. Thorgeirsson (National Cancer Institute, NIH, Bethesda, MD) and is available at the Jackson Laboratory (Stock 016974). *Met*^*EGFP*^ mice were generated as previously described (Kamitakahara et al., 2017; Kast et al., 2017a, 2017b) from GENSAT project clone BX139 (Rockefeller University, RRID:SCR_002721) (Geschwind, 2004). Expression concordance between *Egfp* and *Met* transcript or protein in neurons residing in the neocortex, raphe and vagal motor complex is nearly 100%. While originally generated on mixed backgrounds, both the *Met*^*fx*^ and *Met*^*EGFP*^ transgenic lines were backcrossed for more than 10 generations and maintained isogenically on a C57BL/6J background in our laboratory for all experiments described.

### Method Details

#### Immunohistochemistry

Tissue for immunofluorescence staining was collected on embryonic day (E) 14.5, postnatal day (P) 7, and in P60-P80 (denoted further as P60 or adult) mice. For embryonic tissue collection, timed pregnant breeding pairs were set, with the day of vaginal plug detection designated as E0.5. Sex was not determined for embryonic samples. Embryonic tissue was collected and immersed overnight in fixative (4% paraformaldehyde in 0.1M phosphate-buffered saline (PBS, pH 7.4)). Postnatal mice were deeply anesthetized by intraperitoneal injection of ketamine:xylazine (100mg/kg:10mg/kg, Henry Schein, Melville, NY) and perfused transcardially with 0.9% saline, followed by fixative. Collected tissues were postfixed for two hours, cryoprotected overnight in 20% sucrose in PBS, embedded in Tissue-Tek® Optimal Cutting Temperature Compound and frozen over liquid nitrogen vapors or powdered dry ice. For brain tissue, 20µm-thick cryostat sections were collected in five coronal or sagittal series representing the entire rostral-caudal or medial-lateral extent of the nAmb. For esophageal and laryngeal tissue, 30µm-thick cryostat sections were collected in five coronal series. Slides were stored at -20°C until processed.

For immunofluorescence labeling, slides were incubated for two hours in blocking buffer containing 5% normal donkey serum (Jackson ImmunoResearch, West Grove, PA) and 0.3% Triton X-100 in PBS, then overnight in antibody solution containing 2% normal donkey serum and 0.3% Triton X-100 in PBS with one or more of the following primary antibodies: chicken anti-Green Fluorescent Protein (GFP) (1:500, Abcam Cat# ab13970, RRID:AB_300798), goat anti-MET receptor tyrosine kinase (MET) (1:500, R and D Systems Cat# AF527, RRID:AB_355414), or rabbit anti-Red Fluorescent Protein (RFP) (1:750, Rockland Immunochemicals Cat# 600-401-379). Sections were washed five times for five minutes in PBS, then incubated in antibody solution with one or more of the following secondary or tertiary antibodies: Alexa Fluor® 488 AffiniPure F(ab’)_2_ Fragment Donkey Anti-Chicken IgG (Jackson ImmunoResearch Cat# 703–546-155, RRID:AB_2340376), Biotin-SP-AffiniPure F(ab’)_2_Fragment Donkey Anti-Goat IgG (Jackson ImmunoResearch Cat# 705–066-147, RRID:AB_2340398), Alexa Fluor® 488 Streptavidin (Jackson ImmunoResearch Cat# 016-540-084, RRID: AB_2337249), or Alexa Fluor® 594 AffiniPure F(ab’)_2_ Fragment Donkey Anti-Rabbit IgG (Jackson ImmunoResearch Cat# 711-586-152, RRID:AB_2340622). To label acetylcholine receptor clusters in muscular tissue, α-Bungarotoxin (αBT) Alexa Fluor® 488 conjugate (Thermo Fisher Scientific Cat# B13422) was added to the secondary antibody solution. After several rinses in PBS, sections were counterstained with DAPI (Thermo Fisher Scientific Cat# D1306) and embedded with ProLong Gold Antifade Mountant (Thermo Fisher Scientific Cat# P36930) prior to applying a coverslip.

#### Image Acquisition

A Zeiss LSM 710 laser scanning confocal microscope equipped with 10x, 20x, 40x water-corrected, and 63x oil-corrected objectives was used to acquire immunofluorescence images. Confocal image stacks were collected through the z-axis at a frequency optimally determined by the Zeiss Zen software based on the optics of the microscope and the wavelength of the fluorophores used for analysis. Slides were coded so that the operator was blind to experimental group.

For cell counts of EGFP+ and tdTomato+ neurons in the nAmb, every fifth consecutive coronal section was imaged through the entire rostral-caudal length of the nucleus using a 20x objective. For imaging of laryngeal and esophageal muscle innervation, anatomical regions of interest for each muscle were identified, and one (for each laryngeal muscle) or two (for the esophagus) confocal image stacks were captured using a 20x, 40x water-corrected, or 63x oil-corrected objective.

#### Image Analysis

For cell counts of tdTomato+ and EGFP+ neurons in the nAmb, all image stacks for each animal were manually analyzed using the ‘Cell Counter’ plugin within the Fiji/ImageJ software. For tdTomato+ counts, all neurons that had DAPI colocalized with tdTomato were counted. For EGFP+ counts, all neurons that had DAPI colocalized with both tdTomato and EGFP were counted. Diameters of at least 5% of nuclei were measured in Fiji/ImageJ and averaged per animal. The number of total cells in the nAmb of each animal was estimated in accordance with Abercrombie’s formula (Abercrombie, 1946).

For analyses of laryngeal and esophageal muscle innervation, postsynaptic αBT labeled acetylcholine receptor clusters were counted using the ‘Cell Counter’ plugin in the Fiji/ImageJ software. The number of αBT labeled clusters closely apposed to tdTomato labeled fibers were then counted to generate a measurement of percent innervation. Values from the two esophageal image ROIs were averaged.

#### Ultrasonic Vocalization

Isolation-evoked ultrasonic vocalizations (USVs) were recorded on P7 using a CM16/CMPA ultrasound microphone, positioned 5 cm above the recording chamber floor, and an UltraSoundGate 116H recorder (Avisoft Bioacoustics). Mice were maintained in the home nest environment in a cage warmed by a heating pad until the moment of recording. For each 5-minute recording, a single pup was removed from the nest and recorded in isolation without heat support in the recording chamber. Following each recording, an additional 15-second recording was made to determine whether USVs could be evoked by acute tail pinch. Computed values for USV number and duration were determined using Avisoft-SASLab Pro software. Genotype was unknown to the operator.

Adult male USVs were recorded during a direct social interaction task using a CM16/CMPA ultrasound microphone, positioned 16 cm above the chamber floor, and an UltraSoundGate 116H recorder (Avisoft Bioacoustics). Prior to the day of testing, males from each genotype group were exposed to a 3- to 4-month old C57BL/6J female partner for three days in their home cage to gain social experience. Females were then removed from the cage, and males were housed in isolation for two days to increase motivation to call during the direct social interaction task. On the day of testing, males were habituated to the recording chamber for 10 minutes, then recorded for 6 minutes with a novel 3- to 4-month old C57BL/6J female. Computed values for USV number and duration were determined using Avisoft-SASLab Pro software (Avisoft Bioacoustics, RRID:SCR_014438). Genotype was unknown to the operator.

#### MUPET Analysis

Mice Ultrasonic Profile ExTractor (MUPET) v2.0 (Segbroeck et al., 2017), an open-access MATLAB (MATLAB_R2019a) NeuroResource, was used to analyze USVs recorded on P7 and P60. MUPET uses a complex clustering algorithm to categorize individual USV syllables into a ‘repertoire unit’, based on syllable shape (Segbroeck et al., 2017). The collection of all of the repertoire units made by each genotype or group is referred to as a ‘repertoire’. Audio files were processed in MUPET, and a dataset was created for each genotype or genotype subset. At each age and for each genotype, repertoires of 20 to 140 repertoire units were built to determine the appropriate repertoire size best suited for the analysis. For P7, a repertoire size of 60 units was determined to be optimal based on an overall repertoire modeling score, average log-likelihood, Bayesian Information Criterion, and the number of repertoire units containing only one syllable. The same criteria were used at P60 to determine 100 units as the optimal repertoire size at this age. The ‘Best Match Sorting’ feature was used to generate matrices of Pearson correlations, comparing the similarity of each repertoire unit from the Cre negative dataset to repertoire units in all other genotype or genotype subset datasets. The ‘Unit Activity Sorting’ feature was used to generate a Pearson correlation coefficient for overall repertoire similarity. To estimate the Pearson correlation coefficient for the Cre negative repertoire, six jackknife resampled datasets were analyzed using MUPET (Efron and Stein, 1981; Quenouille, 1956). These jackknife resampled datasets were generated by systematically leaving out one recording in each dataset from the three highest and three lowest vocalizers. Together, the Best Match Sorting and Unit Activity Sorting features measure the similarity of individual repertoire units, and the overall similarity of repertoires between groups, respectively.

#### Quantification and Statistical Analysis

Data were statistically analyzed and graphed using GraphPad Prism software (RRID:SCR_002798), and expressed as mean values ± standard error of the mean. The number of animals required for quantitative analysis was calculated based on power analysis, with the aim of detecting differences between groups that were 1-2 standard deviations from the mean with at least 80% power and p<.05 for significance. Each mouse is considered a sample, with sample sizes of each genotype included in the figure legends for each analysis. For each genotype, a D’Agostino-Pearson normality test was used to determine whether parametric or nonparametric statistical analyses should be performed. For data following a normal distribution, an ordinary one-way ANOVA or a parametric two-tailed unpaired t-test was used to compare means. For data that failed to pass the D’Agostino-Pearson normality test, a nonparametric Kruskal-Wallis test (correcting for multiple comparisons using Dunn’s test) or a nonparametric two-tailed Mann-Whitney test was used to compare mean rank difference. To compare correlation coefficients, Fisher’s r-to-z transformation was applied followed by z-tests with Bonferroni correction. A p<0.05 was used for significance.

## KEY RESOURCES TABLE

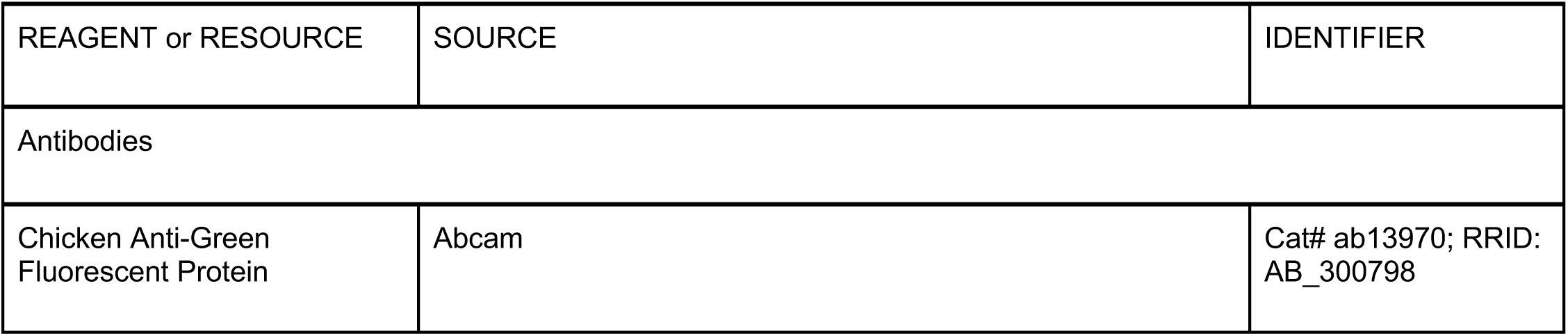
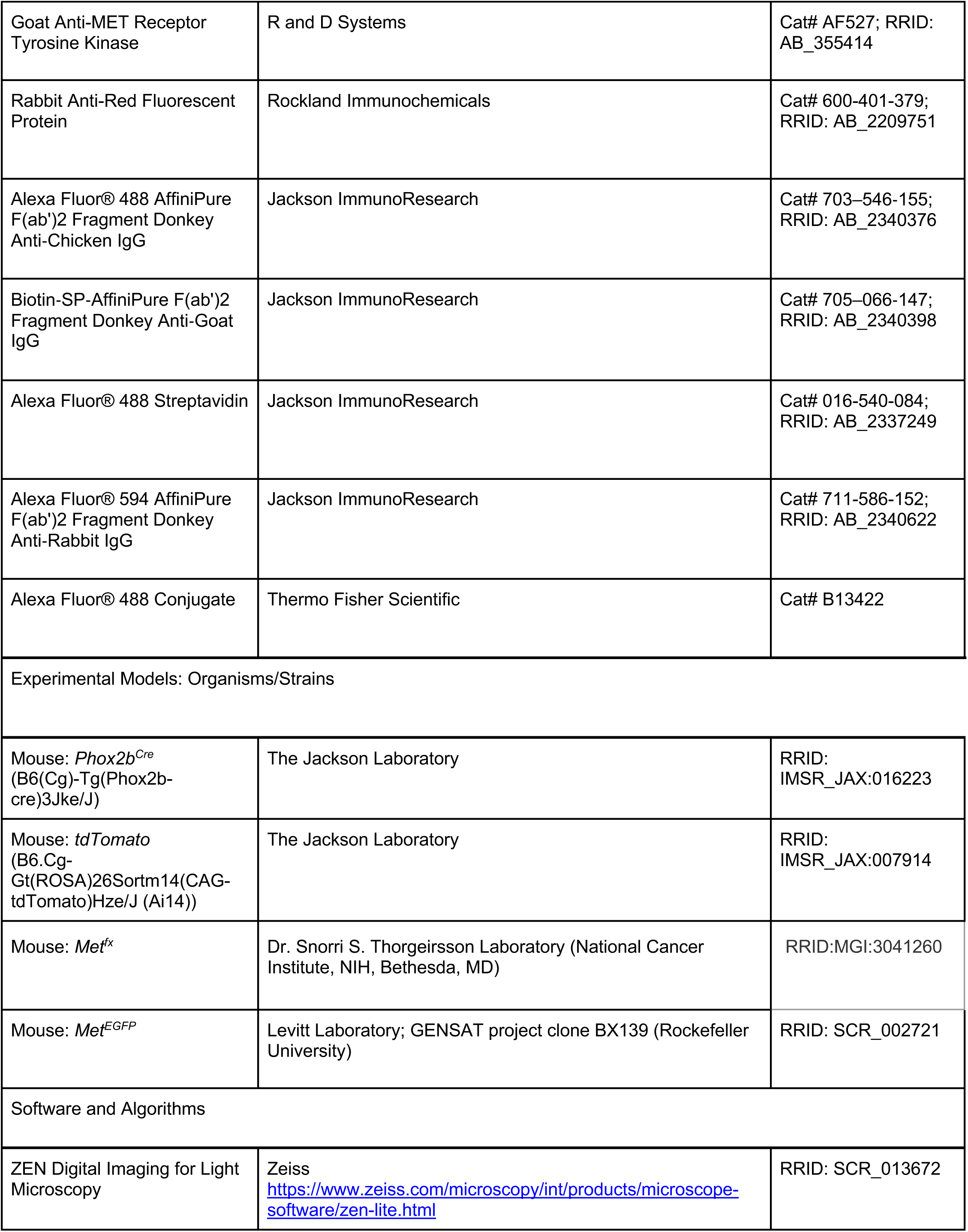
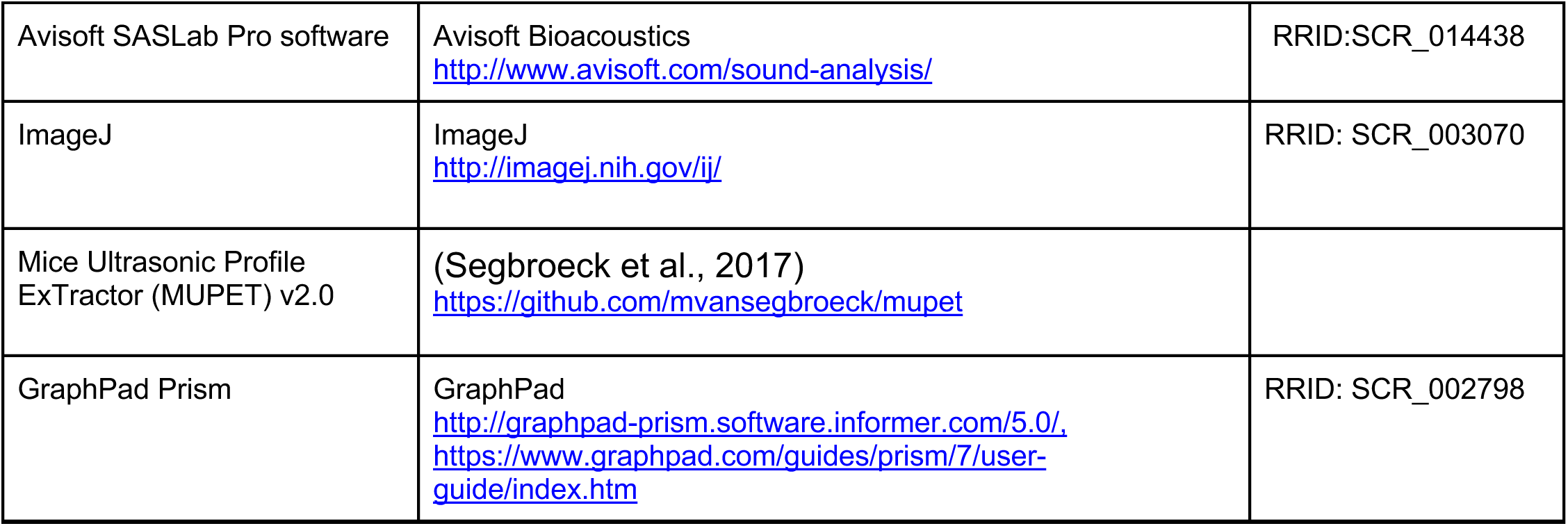

| REAGENT or RESOURCE | SOURCE | IDENTIFIER |
| --- | --- | --- |
| Antibodies |  |  |
| Chicken Anti-Green Fluorescent Protein | Abcam | Cat# ab13970; RRID: AB_300798 |
| Goat Anti-MET Receptor Tyrosine Kinase | R and D Systems | Cat# AF527; RRID: AB_355414 |
| Rabbit Anti-Red Fluorescent Protein | Rockland Immunochemicals | Cat# 600-401-379; RRID: AB_2209751 |
| Alexa Fluor® 488 AffiniPure F(ab')<sub>2</sub> Fragment Donkey Anti-Chicken IgG | Jackson ImmunoResearch | Cat# 703-546-155; RRID: AB_2340376 |
| Biotin-SP-AffiniPure F(ab')<sub>2</sub> Fragment Donkey Anti-Goat IgG | Jackson ImmunoResearch | Cat# 705-066-147; RRID: AB_2340398 |
| Alexa Fluor® 488 Streptavidin | Jackson ImmunoResearch | Cat# 016-540-084; RRID: AB_2337249 |
| Alexa Fluor® 594 AffiniPure F(ab')<sub>2</sub> Fragment Donkey Anti-Rabbit IgG | Jackson ImmunoResearch | Cat# 711-586-152; RRID: AB_2340622 |
| Alexa Fluor® 488 Conjugate | Thermo Fisher Scientific | Cat# B13422 |
| Experimental Models: Organisms/Strains |  |  |
| Mouse: *Phox2b*<sup>Cre</sup> (B6(Cg)-Tg(Phox2b-cre)3Jke/J) | The Jackson Laboratory | RRID: IMSR_JAX:016223 |
| Mouse: *tdTomato* (B6.Cg-Gt(ROSA)26Sortm14(CAG-tdTomato)Hze/J (Ai14)) | The Jackson Laboratory | RRID: IMSR_JAX:007914 |
| Mouse: *Met*<sup>flx</sup> | Dr. Snorri S. Thorgeirsson Laboratory (National Cancer Institute, NIH, Bethesda, MD) | RRID:MGI:3041260 |
| Mouse: *Met*<sup>EGFP</sup> | Levitt Laboratory; GENSAT project clone BX139 (Rockefeller University) | RRID: SCR_002721 |
| Software and Algorithms |  |  |
| ZEN Digital Imaging for Light Microscopy | Zeiss<br>https://www.zeiss.com/microscopy/int/products/microscope-software/zen-lite.html | RRID: SCR_013672 |
| Avisoft SASLab Pro software | Avisoft Bioacoustics<br>http://www.avisoft.com/sound-analysis/ | RRID:SCR_014438 |
| ImageJ | ImageJ<br>http://imagej.nih.gov/ij/ | RRID: SCR_003070 |
| Mice Ultrasonic Profile<br>ExTractor (MUPET) v2.0 | (Segbroeck et al., 2017)<br>https://github.com/mvansegbroeck/mupet |  |
| GraphPad Prism | GraphPad<br>http://graphpad-prism.software.informer.com/5.0/,<br>https://www.graphpad.com/guides/prism/7/user-guide/index.htm | RRID: SCR_002798 |

## Figure Titles and Legends

**Figure S1.**
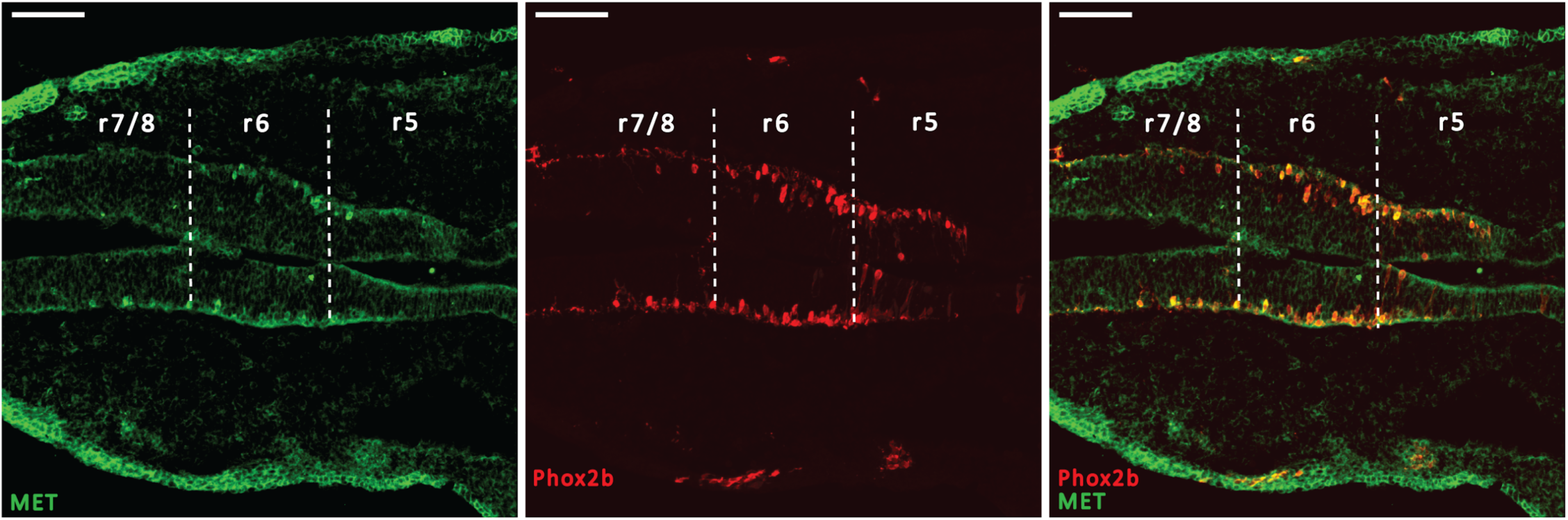
Early Cre-mediated recombination driven by the Phox2b promoter in the developing brainstem. Representative images show MET protein immunoreactivity (green) and Phox2b^cre^;tdTomato endogenous fluorescence (red) on E9.5. r5, rhombomere 5; r6, rhombomere 6; r7/8, rhombomeres 7 and 8. Scale bars = 100μm. The brightness and contrast of each channel was adjusted separately for visualization purposes.

**Figure S2.**
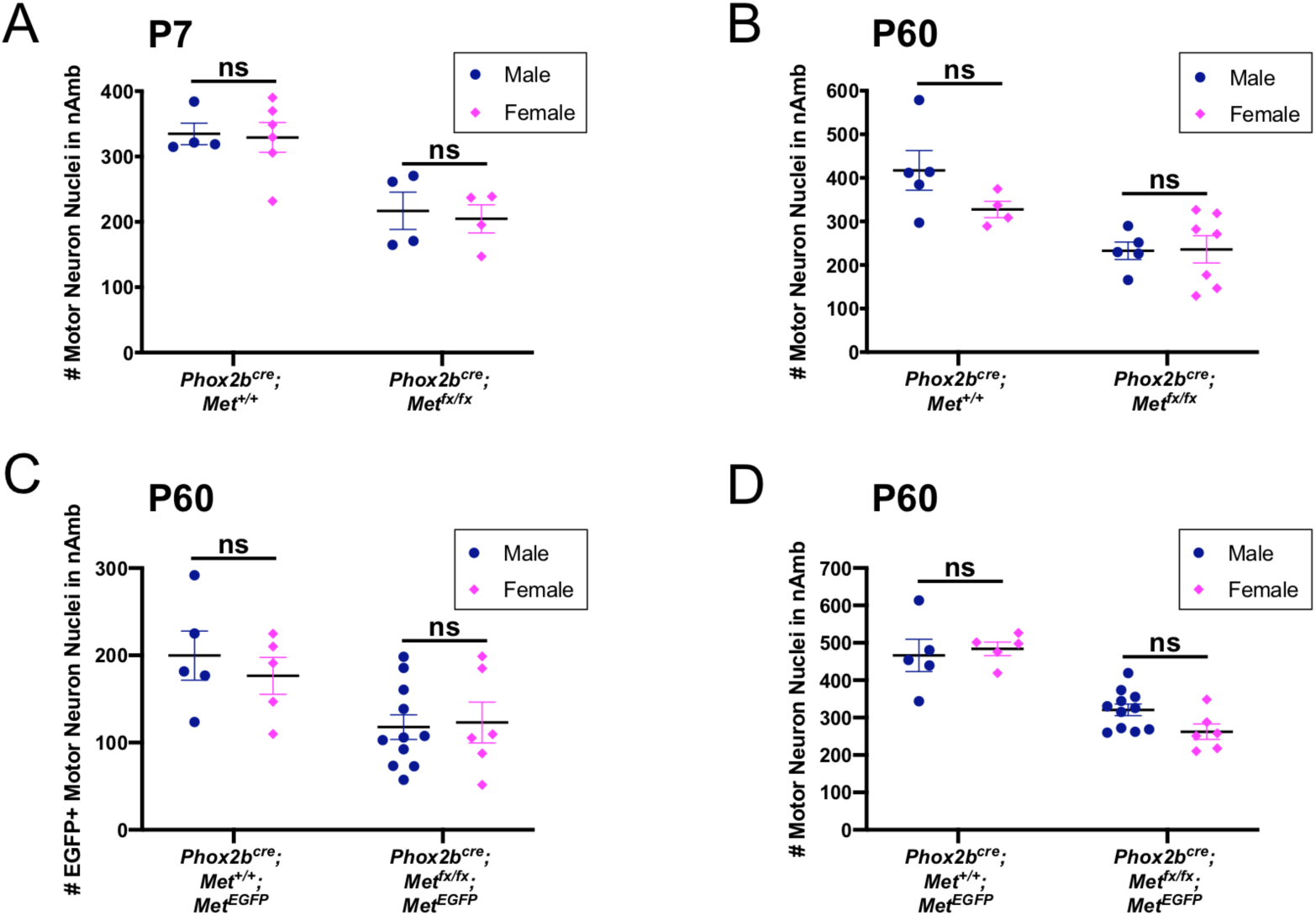
The number of neurons in the nAmb is not different between sexes. (A) tdTom+ cell counts from control and *Phox2b*^*cre*^*;Met*^*fx/fx*^*;tdTom* mice on P7, by sex. n = 4 males, 4 females in the control group; n = 4 males, 4 females in the *Phox2b*^*cre*^*;Met*^*fx/fx*^ group. (B) tdTom+ cell counts from control and *Phox2b*^*cre*^*;Met*^*fx/fx*^*;tdTom* mice on P60, by sex. n = 5 males, 4 females in the control group; n = 5 males, 7 females in the *Phox2b*^*cre*^*;Met*^*fx/fx*^ group. (C) EGFP+ cell counts from control and *Phox2b*^*cre*^*;Met*^*fx/fx*^*;tdTom;Met*^*EGFP*^ mice on P60, by sex. n = 5 males, 5 females in the control group; n = 11 males, 6 females in the *Phox2b*^*cre*^*;Met*^*fx/fx*^ group. (D) tdTom+ cell counts from control and *Phox2b*^*cre*^*;Met*^*fx/fx*^*;tdTom;Met*^*EGFP*^ mice on P60, by sex. n = 5 males, 5 females in the control group; n = 11 males, 6 females in the *Phox2b*^*cre*^*;Met*^*fx/fx*^ group.

